# Emerging fungal pathogen on an invasive grass differentially affects native species

**DOI:** 10.1101/2020.08.06.239319

**Authors:** Amy E. Kendig, Vida J. Svahnstrom, Ashish Adhikari, Philip F. Harmon, S. Luke Flory

**Author notes:** These authors contributed equally to this work (Amy E. Kendig, Vida J. Svahnstrom).

## Abstract

Infectious diseases and invasive species are strong drivers of biological systems that may interact to shift plant community composition. Disease and invasion can each directly suppress native populations, but variation in responses among native species to disease, invasion, and their combined effects are not well characterized. Here, we quantified the responses of three native North American grass species to experimental inoculation with the fungal pathogen *Bipolaris gigantea*, which has recently emerged in populations of the invasive grass *Microstegium vimineum*, causing leaf spot disease. In a greenhouse experiment, we examined the direct effects of disease on the native species and the indirect effects of disease on the native species through altered competition with *M. vimineum*, which was planted at a range of densities. Pathogen inoculation directly affected each of the three native species in unique ways, by increasing, decreasing, or not changing their biomass relative to mock inoculation. Higher *M. vimineum* densities, however, reduced the biomass of all three native species, regardless of inoculation treatment, suggesting that disease had no indirect effects through altered competition. In addition, competition with *M. vimineum* suppressed native plant biomass to a greater extent than disease. The differential impacts of *B. gigantea* and the consistent impacts of *M. vimineum* on native species biomass suggest that disease may modify native plant community composition while plant invasion may suppress multiple native plant species in systems where these drivers co-occur.

## Introduction

Both plant invasions and infectious diseases can afflict natural plant communities [1,2] by reducing plant diversity and biomass production [3–5]. Invasive species and disease outbreaks can co-occur in communities because the species are co-introduced, or because invasive species amplify disease transmission [6]. The combined effects of plant invasion and disease can impact communities in a number of ways: by maintaining community structure, by suppressing native species more than would either driver on its own, or by slowing or even reversing the invasion [7,8]. Which of these outcomes occurs depends on the responses of the invasive species and the co-occurring native species to infection [9,10]. Pairwise tests of disease-mediated competition between native and invasive plant species have identified possible outcomes of invasion and disease [11–14]. However, the relevant guild of native species in natural communities is often diverse, and it is unclear how native plant species vary in their responses to the combined effects of invasion and disease.

Disease can affect native species in at least two ways: by directly infecting native species and suppressing their survival or performance [5], or indirectly by modifying competition with other species [15]. Invasive species may enhance the transmission of pathogens or the disease impacts experienced by native species, which is termed “spillover” when the pathogen originates from the invasive species and “spillback” when the pathogen persists locally in the absence of the invasive species [16,17]. For example, high densities of invasive cheatgrass (*Bromus tectorum*) promoted spillover of a fungal pathogen to native plant seeds [18]. Pathogen infection of native plants can reduce growth, reproduction, and survival [5], as well as induce compensatory growth and reproduction [19,20]. The impacts of disease on natural plant populations and communities can, at times, match or exceed those of herbivores [21].

Disease can promote coexistence or competitive exclusion between invasive and native species [15]. If disease disproportionately impacts the invasive species, reductions in growth and resource uptake may release the native species from competition, which occurred when a powdery mildew fungus infected garlic mustard (*Alliaria petiolata*) [14]. In contrast, disease-induced fitness costs may reduce native species’ competitive ability, which is hypothesized to have promoted invasion of European grasses in California [11,13]. Plant species can vary widely in their competitive ability when interacting with invasive plants [22,23], their susceptibility to infection [24,25], and their performance losses due to disease [20,26]. Differential responses of native species to invasion and disease could determine how the native plant community responds to disease-mediated competition.

Here we investigated how a pathogen that is known to suppress growth and reproduction of an invasive plant species [24,27] might directly or indirectly affect three native plant species. First, we evaluated the direct effects of pathogen inoculation on the invasive and native species in a greenhouse experiment, hypothesizing that disease would suppress invasive species growth [24,27]. We were uncertain about how disease would affect the native species, but acknowledged that a range of outcomes were possible given interspecific variation in host-pathogen interactions [24–26]. Then, we examined whether pathogen inoculation modified the competitive effect of the invasive species on each of the native species using an additive competition experimental design [28,29]. We hypothesized that pathogen inoculation would reduce the competitive effect of the invasive species, but that disease-induced losses experienced by some native species would exacerbate the impacts of invader competition.

## Materials and methods

### Study system

*Microstegium vimineum* (stiltgrass, hereafter *Microstegium*) is an annual grass species native to Asia that was first recorded in the wild in the United States in 1919 [30]. *Microstegium* forms dense populations and litter layers in eastern and midwestern U.S. forest understories, suppressing herbaceous plants and tree seedlings [23,31]. Over the last two decades, *Microstegium* has acquired fungal leaf spot infections from species in the genus *Bipolaris* that reduce its biomass and seed production [24,27]. *Dichanthelium clandestinum* (deer-tongue grass, hereafter *Dichanthelium*), *Elymus virginicus* (Virginia wild rye, hereafter *Elymus*), and *Eragrostis spectabilis* (purple lovegrass, hereafter *Eragrostis*) are grass species that are native to the U.S., co-occur with *Microstegium*, and may be susceptible to infection by *Bipolaris* spp. that infect *Microstegium* [24,27,32,33].

We collected *Microstegium* seeds from Big Oaks National Wildlife Refuge (BONWR) in Madison, IN, USA in 2015. We obtained *Elymus* and *Eragrostis* seeds from Prairie Moon Nursery (Winona, MN, USA) and *Dichanthelium* seeds from Sheffield’s Seed Company (Locke, NY, USA) in 2018. All seeds were stored at 4°C. The fungal isolate of *Bipolaris gigantea* originated from a litter competition study conducted in a greenhouse in Gainesville, FL, USA (S1 Appendix). Briefly, dried *Microstegium* litter that had been collected from BONWR in 2018 was placed in a layer over pots of *Microstegium* seedlings, which were enclosed with thin transparent plastic bag to maintain high humidity. Seedlings developed eyespot lesions with light centers and dark brown edges, characteristic of *Bipolaris* infection (Fig. 1C). Leaves with eyespot lesions were collected and incubated at 28°C for 24 h. Large conidia on conidiophores characteristic of *B. gigantea* [34] were observed in lesions and were transferred by sterilized dissecting needle to half-strength V8 media agar plates. Hyphal tip transfers of the single spore colony to additional agar plates were made to obtain a pure fungal culture, BGLMS-1. The isolate was stored on sterile filter paper at 4°C [27].

**Fig. 1.**
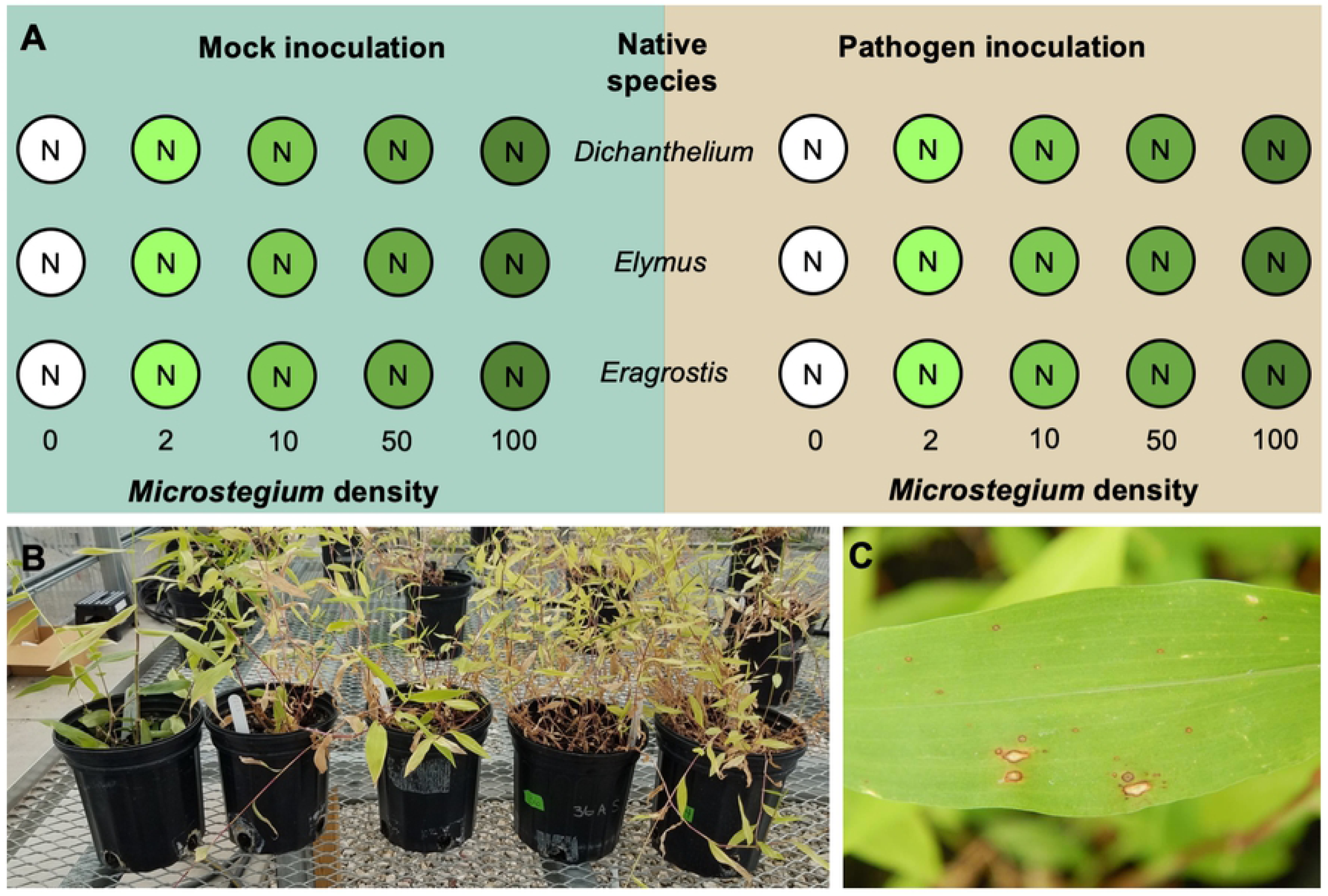
Experimental methods. Illustration of (A) the experimental design, (B) an example of the realized *Microstegium* density gradient (with *Dichanthelium* as the native species), and (C) an example of *Bipolaris*-like lesions on a *Microstegium* leaf from the experiment. Circles in A represent 1 L pots, with “N” indicating the central position of the native plant and the intensity of green shading indicating the *Microstegium* density (planted density values labelled below pots). Each represented pot was replicated four times.

### Greenhouse experiment

We performed the experiment in a greenhouse in Gainesville, FL, USA, from June 26, 2019 to September 12, 2019. The potting mix used in the experiment (Jolly Gardener Pro-Line Custom Growing Mix) was autoclaved at 120–130°C for 30 minutes and all pots and trays were sprayed with 10% bleach solution (0.6% sodium hypochlorite) and rinsed with tap water after approximately five minutes to minimize risk of contamination by non-focal pathogens. To quantify the effect of *Microstegium* competition on the native species, we planted one individual of a native species in the center of each 1 L pot and 0, 2, 10, 50, or 100 *Microstegium* around the native plant [28,29] (Fig. 1A). The native species were transplanted from germination trays to the 1 L pots after growing in the greenhouse for 21 days, and the *Microstegium* were added to the 1 L pots as seeds (50 and 100 seed numbers estimated by weight). We chose native plant individuals that were similar in size (2 to 3 true leaves) to transplant into pots for the experiment. The 15 plant combinations (each of the three native species with five *Microstegium* densities) were replicated eight times, half of which were inoculated with *B. gigantea* and half of which were mock-inoculated.

To prepare inoculum for the experiment, the *B. gigantea* isolate (BGLMS-1) was revived from 4°C storage by placing colonized, 3 to 5 mm diameter, filter paper pieces on half-strength V8 media agar plates. Fungal colonies grew under 12 h day and night fluorescent light at 26°C for one week and were transferred to new half-strength V8 media agar plates. Conidia were harvested from fungal colonies by flooding plates with 10 ml of sterile deionized water with 0.1% Tween 20 (Sigma-Aldrich, St. Louis, MO, USA). The resulting conidia suspension was filtered through a layer of cheese cloth, and conidia were quantified with a Spencer Bright-Line hemacytometer (American Optical Company, Buffalo, NY, USA). The concentration of inoculum was adjusted to 15,000 conidia/ml and applied to plants with a Passche H-202S airbrush sprayer (Kenosha, WI, USA). Inoculations occurred six days after planting, and half of the pots were sprayed until runoff with the spore suspension while the other half were sprayed with the same volume of water with 0.1% Tween 20 (i.e., mock inoculation control). To encourage infection, we placed a wet paper towel in each pot and sealed each pot with a transparent plastic bag secured with a rubber band. The plastic bags and paper towels were removed after seven days [35].

Prior to inoculation, plants were hand-watered and then watered with a lawn sprinkler daily, during which time we constructed a drip irrigation system. Following inoculation, the plants were watered daily for eight minutes with a drip irrigation system to help control for variation in water application. Ten days after bag removal, all plants were sprayed with Garden Safe insecticidal soap (Bridgeton, MO, USA) to help control aphids and thrips.

### Data collection

To assess disease incidence 14 days after inoculation, we recorded the number of *Microstegium* leaves with *Bipolaris*-like lesions (Fig. 1C) and the total number of leaves for three *Microstegium* plants (or two plants for the pots with only two) per pot. The number of leaves per plant were averaged within pots and multiplied by the total number of plants per pot, based on seeding rate, to estimate the total number of *Microstegium* leaves per pot. We counted the number of leaves with lesions and the total number of leaves of native plants that received the pathogen inoculation treatment and had lesions. None of the plants in mock-inoculated pots had lesions with one exception: in one pot that contained *Dichanthelium* and 100 *Microstegium* plants, 46 *Microstegium* leaves had lesions. We concluded that this pot was inadvertently inoculated and removed it from the analyses. To assess plant performance, we harvested the aboveground biomass of all pots on September 12, 2019, separated the native plants from the *Microstegium*, dried the biomass at 60°C to constant mass, and weighed it.

### Statistical analyses

To evaluate disease incidence experienced by plants across the *Microstegium* density gradient, we fit a generalized linear regression to the estimated proportion of *Microstegium* leaves with lesions per pathogen-inoculated pot using *Microstegium* density (the number of *Microstegium* seeds added to each pot), native species identity, and their interaction as the explanatory variables. The model was fit with Bayesian statistical inference using the brm function in the brms package [36], an interface for Stan [37], in R version 3.5.2 [38]. The model contained three Markov chains with 6000 iterations each and a discarded burn-in period of 1000 iterations. We chose prior distributions based on whether model variables could reasonably take on negative values in addition to positive values (Gaussian or Cauchy) or not (gamma or exponential). We chose parameters for prior distributions that reflected limited a priori information about variable values. We used a binomial response distribution and a Gaussian distribution for the intercept and coefficient priors (location = 0, scale = 10). There were too few native plant leaves with lesions to statistically analyze disease incidence, nevertheless, we present these results graphically to assess qualitative patterns.

To evaluate the effects of *Microstegium* density and pathogen inoculation on *Microstegium* performance, we fit a linear regression to *Microstegium* biomass:

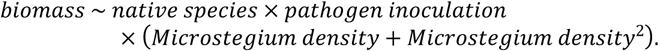

This formulation allowed us to estimate separate quadratic relationships between *Microstegium* biomass and *Microstegium* density for each native species–inoculation treatment combination. We used a Gaussian response distribution, a Gaussian distribution for the intercept prior (location = 2, scale = 10) and the coefficient priors (location = 0, scale = 10), and a Cauchy distribution for the standard deviation prior (location = 0, scale = 1). Otherwise, the model was fit using the same methods described for disease incidence.

To evaluate the direct effects, and indirect effects through *Microstegium* competition, of inoculation treatment on native plant performance, we fit a Beverton-Holt function to native plant biomass:

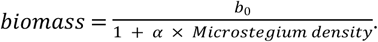

We fit this function to all of the native plant biomass data, estimating separate b_0_ (biomass in the absence of competition) and α (the competitive effect of *Microstegium*) values for each native species–inoculation treatment combination. We used a Gaussian response distribution, a Gamma distribution for the b_0_ prior (shape = 2, scale = 1), an exponential distribution for the α prior (rate = 0.5), and a Cauchy distribution for the standard deviation prior (location = 0, scale = 1). Otherwise, the model was fit using the same methods described for disease incidence. To evaluate differences in b_0_ and α between treatments, we subtracted the estimate for one treatment from the other for each posterior sample (*n* = 1500) and then calculated the mean and 95% highest posterior density interval using the mean_hdi function in the tidybayes package [39]. To assess model fits, we checked that the r-hat value for each parameter was equal to one, visually examined convergence of the three chains, and compared the observed data to simulated data from the posterior predictive distributions using the pp_check function [36]. We report results as statistically significant if their 95% credible intervals omit zero. We used the tidyverse packages to clean data and create figures [40].

## Results

Disease incidence on *Microstegium* decreased with increasing *Microstegium* density, especially when *Microstegium* was grown with *Dichanthelium* and *Eragrostis* (S1 Table). This pattern arose because pots with higher *Microstegium* densities had more leaves (despite smaller plants, S1 Fig.), but similar numbers of leaves with lesions as pots with lower *Microstegium* densities (Fig. 2).

**Fig. 2.**
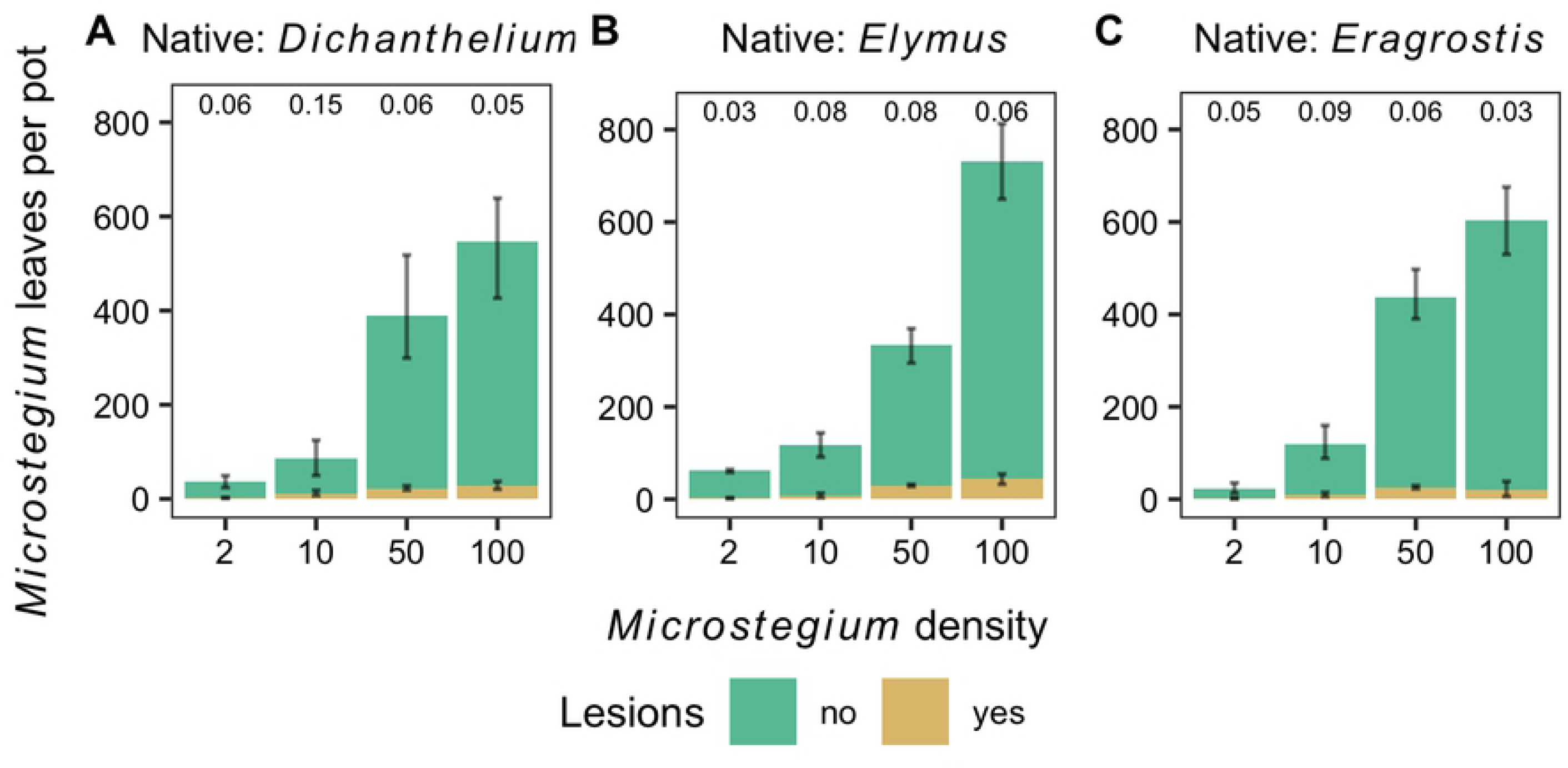
*Microstegium* disease incidence. The effect of *Microstegium* density (seeds per pot) on the estimated proportion of *Microstegium* leaves with lesions following pathogen inoculation in the presence of (A) *Dichanthelium*, (B) *Elymus*, and (C) *Eragrostis* (mean ± 95% confidence intervals). All leaves with lesions were counted and the total number leaves per pot were estimated by counting the number of leaves on up to three plants per pot. The average estimated proportion of leaves with lesions for each density level are printed along the top of each panel.

Pathogen inoculation resulted in lesions on all three native plant species but only in the presence of *Microstegium* (Fig. 3). *Elymus* was the most susceptible native species, with lesions forming on seven out of 20 plants (Fig. 3B), relative to three out of 20 plants for each of the other species (Fig. 3A, C). Of the plants with lesions, higher *Microstegium* density tended to increase the proportion of leaves with lesions, for example, 17% of *Dichanthelium* leaves had lesions when grown with 100 *Microstegium* vs. 10% with 10 *Microstegium*. Similarly, 38% of *Elymus* leaves had lesions when grown with 100 *Microstegium* vs. 23% with 10 *Microstegium*.

**Fig. 3.**
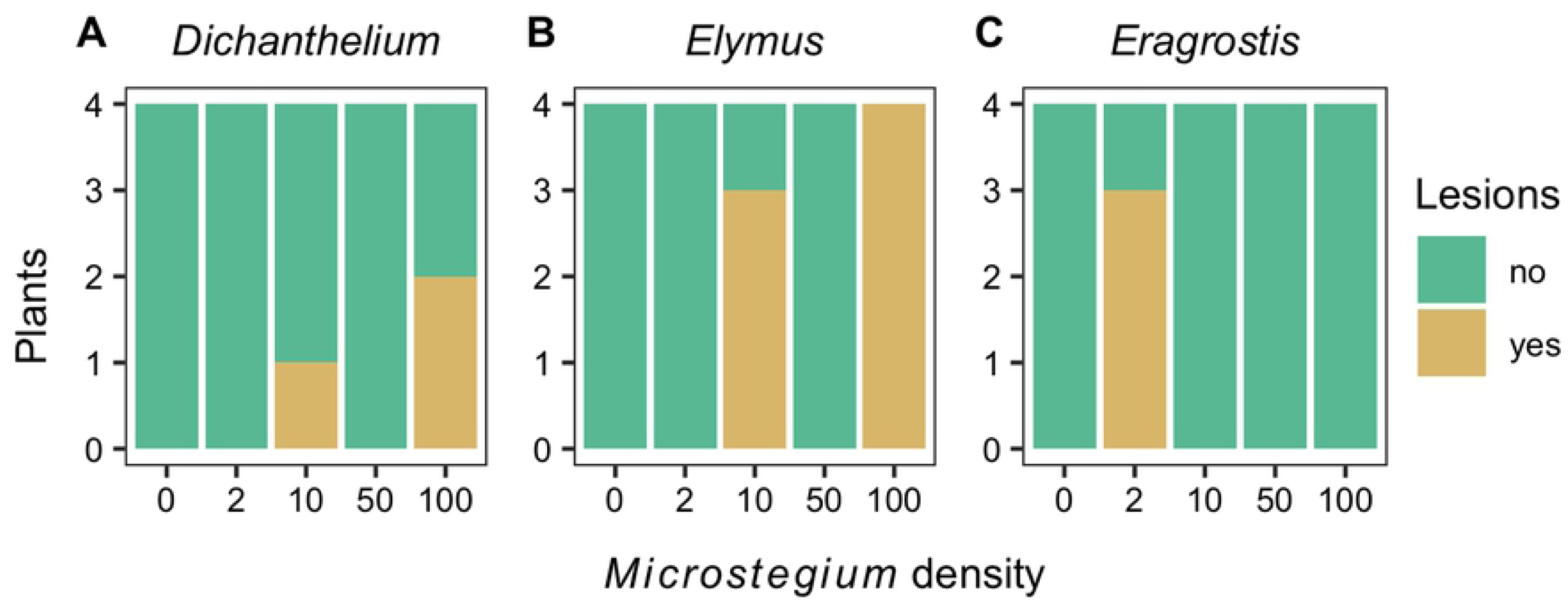
Native plant disease incidence. The effect of *Microstegium* density (seeds per pot) on the number of (A) *Dichanthelium*, (B) *Elymus*, and (C) *Eragrostis* plants with lesions following pathogen inoculation.

Pathogen inoculation did not alter *Microstegium* biomass relative to the mock inoculation control (Fig. 4, S2 Table). *Microstegium* biomass increased with *Microstegium* density (S2 Table) and tended to vary in shape, although not significantly, when grown with the three species: saturating at high densities when grown with *Dichanthelium* (Fig. 4A), increasing nearly linearly when grown with *Elymus* (Fig. 4B), and peaking at intermediate densities when grown with *Eragrostis* (Fig. 4B).

**Fig. 4.**
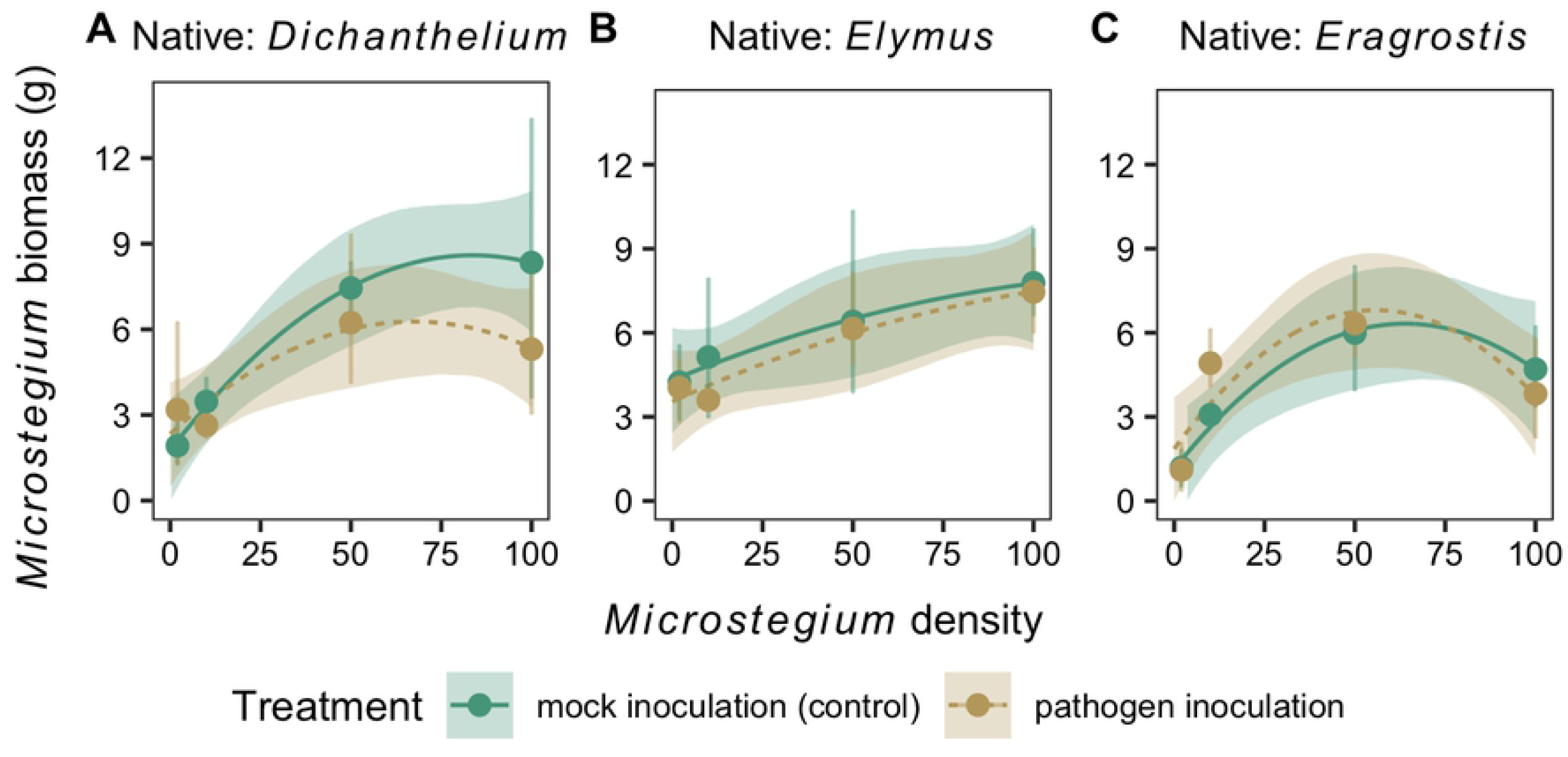
*Microstegium* biomass. The effect of *Microstegium* density (seeds per pot) and inoculation treatment on the biomass of *Microstegium* in the presence of (A) *Dichanthelium*, (B) *Elymus*, and (C) *Eragrostis*. Observations (points and error bars, mean ± 95% confidence intervals) and model fits (lines and shaded ribbons, mean ± 95% credible intervals) are shown.

The direct effects of pathogen inoculation on native plant biomass depended on the native plant species (S3 Table). In the absence of *Microstegium*, pathogen inoculation increased *Dichanthelium* biomass (Fig. 5A), decreased *Elymus* biomass (Fig. 5B), and had no effect on *Eragrostis* biomass (Fig. 5C) relative to the mock inoculation control. The effect of *Microstegium* density on native biomass was consistent across the native species (S3 Table) and was not modified by the inoculation treatment (Fig. 5).

**Fig. 5.**
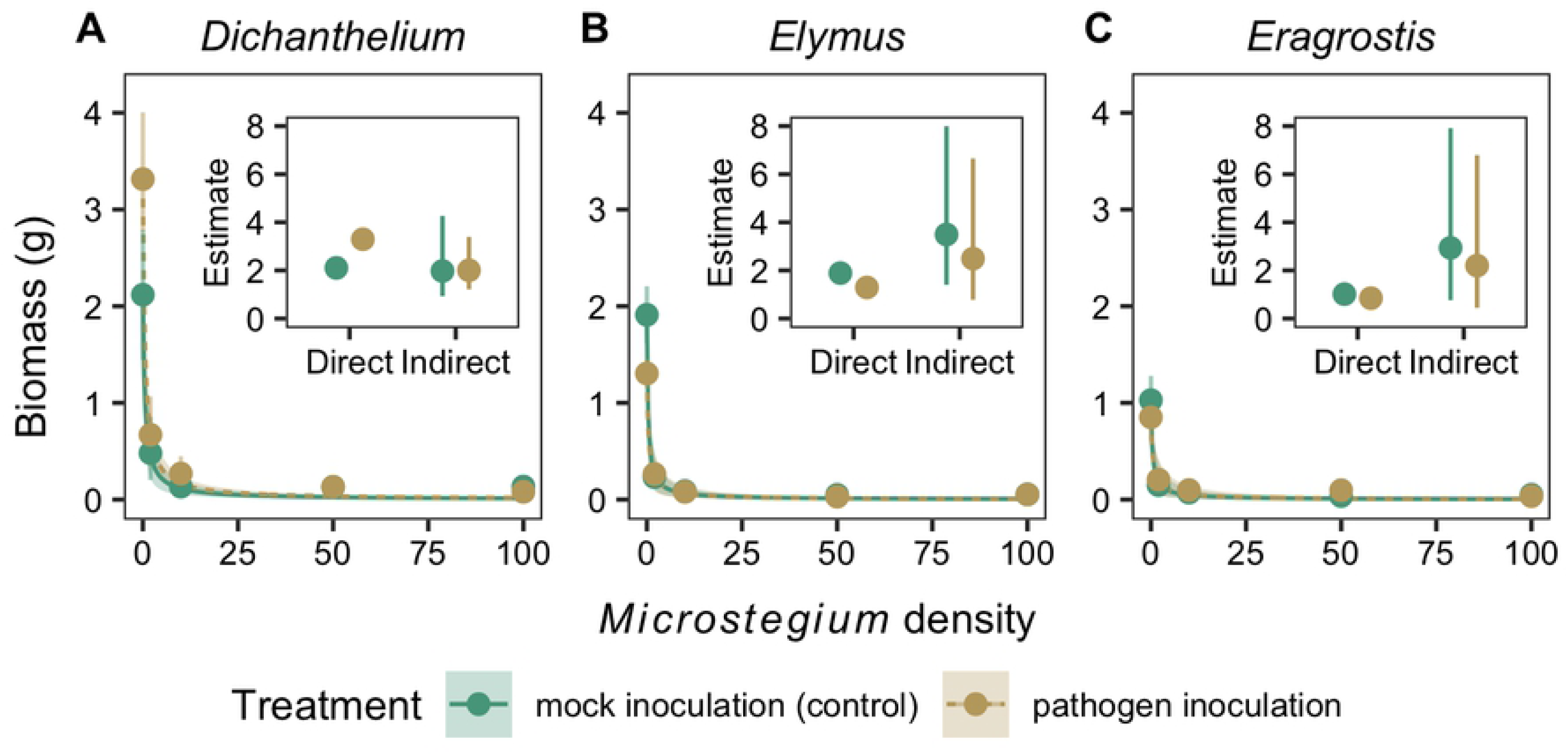
Native plant biomass. The effect of *Microstegium* density (seeds per pot) and inoculation treatment on the biomass of (A) *Dichanthelium*, (B) *Elymus*, and (C) *Eragrostis*. Main plots show observations (points and error bars, mean ± 95% confidence intervals) and model fits (lines and shaded ribbons, mean ± 95% credible intervals). Inset plots show the model-estimated direct (b_0_) and indirect (α) effects of the inoculation treatment on native plant biomass (mean ± 95% credible intervals).

## Discussion

We evaluated how a pathogen that is known to suppress the performance of an invasive plant [24,27] directly, and indirectly via modification of competition, affected three native species. Pathogen inoculation did not significantly affect *Microstegium* biomass relative to the mock inoculation control and it had contrasting effects on the native plant species by increasing *Dichanthelium* biomass, decreasing *Elymus* biomass, and having no effect on *Eragrostis* biomass. The negative effect of *Microstegium* density on biomass for each of the native species was the same whether plants were inoculated with the pathogen or the control. These results suggest that disease caused by *Bipolaris* may alter the community composition of native species, but that *Microstegium* invasion has a consistently large negative effect on native species, potentially overshadowing the effects of disease.

Experimentally suppressing *Bipolaris* infection using fungicide in the field increased *Microstegium* biomass by 33–39% [24,27], suggesting substantial effects of severe disease symptom development. However, despite using a pathogenic *Bipolaris* isolate in our experiment, inoculation caused relatively low levels of disease incidence (relative to approximately 40% of leaves with lesions documented in the field site where infected litter was collected, Kendig et al. unpublished data), which had no effect on *Microstegium* biomass relative to the mock inoculation control. Plant disease transmission and incidence depends on the favorability and duration of environmental conditions and the inoculum load [41,42]. Our experiment relied on a single inoculation and extended incubation; however, field conditions that result in cycles of leaf wetness events (e.g., dew or precipitation) can enhance fungal infection [41] and promote multiple disease cycles throughout the growing season. The concentration of *Bipolaris* conidia in our experimental inoculations was limited by the number of conidia we could harvest from agar plates in the lab and was relatively low (15,000 conidia/ml compared to e.g., 10^5^ conidia/ml [42]). While the conditions for leaf wetness and conidia suspension concentration likely limited the possible extent of disease incidence, they may reflect initial disease dynamics in the field, which is consistent with the age of plants we used in the experiment. Because inoculated *Microstegium* plants had lesions and plants in the control treatment did not, it is also possible that similar biomass responses between plants in each of the inoculation treatments was due to tolerance or compensatory growth during the initial cycle of disease progression. In contrast, the absence of a pathogen inoculation effect on *Eragrostis*, which had few lesions, may be because of disease resistance.

It is likely that spillover or spillback of pathogens from invasive to native plants will have distinct effects on different native species [18,26]. The three native species in the experiment demonstrated unique responses to pathogen inoculation in the absence of *Microstegium*.

Interestingly, no fungal lesions were observed on these native plants despite seven days of incubation inside plastic bags. The absence of visible symptoms, however, does not necessarily indicate a lack of infection. For example, some fungi are asymptomatic endophytes of invasive Crofton weed (*Ageratina adenophora*) but cause visible leaf spots on co-occurring plant species [43]. Future efforts that aim to better characterize host-pathogen interactions between *Microstegium* or native plant species and *Bipolaris* fungi could confirm infection by attempting to re-isolate the fungus after disease symptoms do or don’t develop. The range of pathogen inoculation effects, from negative (on *Elymus*) to positive (on *Dichanthelium*), is consistent with variation in *Bipolaris* disease incidence across species [33] and adheres to the theory of a context-dependent mutualism-parasitism continuum that has been applied to microbes such as mycorrhizae [44]. Plant-soil feedback studies suggest that the relative abundance of species in the field can be predicted by growth responses to soil microbe manipulations [45,46]. However, whether disease-induced changes in growth, survival, or reproduction of native and invasive plant species are sufficient to plant shift community structure is an important area of research [15].

The available studies on disease-mediated competition between invasive and native plant species indicate that infection of invasive plants can contribute to either native plant persistence or recovery [12,14,27,47]. However, in our experiment, inoculation with *Bipolaris* did not modify the effect of *Microstegium* density on the three native species relative to the mock inoculation control, suggesting that competitive effects of *Microstegium* on native species are likely to be consistent in the presence or absence of low levels of disease incidence. The magnitude of competitive effects of *Microstegium* on native plant biomass under these conditions far exceeded those of pathogen inoculation, which may be common across plant communities [48]. Because our experimental methods and conditions may have limited the impacts of *Bipolaris* leaf spot disease, it is crucial to explore disease-mediated competitive effects of *Microstegium* in the field or with methods that may result in disease incidence approaching levels observed in the field. Nonetheless, the impacts of *Bipolaris* on *Microstegium* competition may simply be minor, as has been demonstrated for disease effects on cheatgrass competition [12] and herbivory effects on Amur honeysuckle (*Lonicera maackii*) competition with native species [49]. In that case, competitive effects of *Microstegium* on native species are likely to overshadow the effects of *Bipolaris*-induced disease.

We used a greenhouse experiment to demonstrate that a fungal leaf spot pathogen that has accumulated on a widespread invasive grass has differential effects on native species but does not modify native–invasive plant competition. Complementary experiments in the field could help determine whether these findings are consistent when disease impacts on the invasive species are stronger. Transmission of *Bipolaris* may depend on *Microstegium* densities, potentially creating feedbacks between infection and density, which we controlled for in our experiment. The competitive effects of native plant species on *Microstegium* in the presence and absence of disease also may be important for understanding long-term community dynamics [8]. The emergence of infectious diseases in invaded plant communities may lead to natural biological control of the invasive species [7], exacerbated effects of invasion if the pathogen spills over or spills back to native species [11], or there may be no effect of disease [8]. Altogether, our study suggests that the presence of disease in this system is unlikely to modify the large negative impact of the invasive species on native species.

## Acknowledgements

We would like to thank Liliana Benitez, Zobia Chanda, Laney Davidson, Zadok Jollie, and Shannon Regan for their assistance with the experiment, Simon Riley for his assistance with the statistical analysis, and Brett Lane, Erica Goss, Robert Holt, Michael Barfield, Nicholas Kortessis, Margaret Simon, Christopher Wojan, and Keith Clay for discussions about the manuscript.

## Supporting Information Captions

**S1 Appendix. Extended methods of *Bipolaris gigantea* isolate collection**.

**S1 Table. *Microstegium* disease incidence model**. Model-estimated parameters for generalized linear regression of the estimated proportion of *Microstegium* leaves with lesions from the pathogen inoculation treatment. Estimate is the mean and Est. Error is the standard deviation of the posterior distribution. Q2.5 and Q97.5 are the lower and upper 95% credible intervals, respectively. Estimates with 95% credible intervals that exclude zero are in bold.

**S2 Table. *Microstegium* biomass model**. Model-estimated parameters for linear regression model of *Microstegium* biomass. Estimate is the mean and Est. Error is the standard deviation of the posterior distribution. Q2.5 and Q97.5 are the lower and upper 95% credible intervals, respectively. Estimates with 95% credible intervals that exclude zero are in bold.

**S3 Table. Native plant biomass model**. Model-estimated direct and indirect effects of pathogen inoculation on native plant biomass for each native species and competitive effects of *Microstegium* on the native species under mock inoculation. Estimate is the mean and Q2.5 and Q97.5 are the lower and upper 95% credible intervals, respectively. Estimates with 95% credible intervals that exclude zero are in bold.

**S1 Fig. Estimated *Microstegium* leaves**. The effect of *Microstegium* density on the estimated number of *Microstegium* leaves per plant when grown in the presence of (A) *Dichanthelium*, (B) *Elymus*, and (C) *Eragrostis* (mean ± 95% confidence intervals). Up to three plants were counted per pot.

## References

1. Rejmánek M, Richardson DM, Pyšek P. Plant invasions and invasibility of plant communities. In: van der Maarel E, editor. Vegetation Ecology. Malden, MA, USA: Blackwell Science Ltd; 2005. pp. 332–355.

2. Alexander HM. Disease in natural plant populations, communities, and ecosystems: Insights into ecological and evolutionary processes. Plant Dis. 2010;94: 492–503. doi: 10.1094/PDIS-94-5-0492

3. Cappelli SL, Pichon NA, Kempel A, Allan E. Sick plants in grassland communities: a growth-defense trade-off is the main driver of fungal pathogen abundance. Ecol Lett. 2020; 806299. doi: 10.1101/806299

4. Vilà M, Espinar JL, Hejda M, Hulme PE, Jarošík V, Maron JL, et al. Ecological impacts of invasive alien plants: A meta-analysis of their effects on species, communities and ecosystems. Ecol Lett. 2011;14: 702–708. doi: 10.1111/j.1461-0248.2011.01628.x

5. Dobson A, Crawley M. Pathogens and the structure of plant communities. Trends Ecol Evol. 1994;9: 393–398. doi: 10.1016/0169-5347(94)90062-0

6. Dunn AM, Hatcher MJ. Parasites and biological invasions: Parallels, interactions, and control. Trends Parasitol. 2015;31: 189–199. doi: 10.1016/j.pt.2014.12.003

7. Hilker FM, Lewis MA, Seno H, Langlais M, Malchow H. Pathogens can slow down or reverse invasion fronts of their hosts. Biol Invasions. 2005;7: 817–832. doi: 10.1007/s10530-005-5215-9

8. Flory SL, Clay K. Pathogen accumulation and long-term dynamics of plant invasions. J Ecol. 2013;101: 607–613. doi: 10.1111/1365-2745.12078

9. Holt RD, Grover J, Tilman D. Simple rules for interspecific dominance in systems with exploitative and apparent competition. Am Nat. 1994;144: 741–771. doi: 10.1086/285705

10. Mordecai EA. Consequences of pathogen spillover for cheatgrass-invaded grasslands: coexistence, competitive exclusion, or priority effects. Am Nat. 2013;181: 737–747. doi: 10.1086/670190

11. Eppinga MB, Rietkerk M, Dekker SC, De Ruiter PC, Van Der Putten WH. Accumulation of local pathogens: A new hypothesis to explain exotic plant invasions. Oikos. 2006. pp. 168–176. doi: 10.1111/j.2006.0030-1299.14625.x

12. Mordecai EA. Despite spillover, a shared pathogen promotes native plant persistence in a cheatgrass-invaded grassland. Ecology. 2013;94: 2744–53. doi: 10.1890/13-0086.1

13. Borer ET, Hosseini PR, Seabloom EW, Dobson AP. Pathogen-induced reversal of native dominance in a grassland community. PNAS. 2007;104: 5473–8. doi: 10.1073/pnas.0608573104

14. Cipollini D, Enright S. A powdery mildew fungus levels the playing field for garlic mustard (Alliaria petiolata) and a North American native plant. Invasive Plant Sci Manag. 2009;2: 253–259. doi: 10.1614/IPSM-08-144.1

15. Mordecai EA. Pathogen impacts on plant communities: unifying theory, concepts, and empirical work. Ecol Monogr. 2011;81: 429–441. doi: 10.1890/10-2241.1

16. Power AG, Mitchell CE. Pathogen spillover in disease epidemics. Am Nat. 2004;164: S79–S89. doi: 10.1086/424610

17. Kelly DW, Paterson RA, Townsend CR, Poulin R, Tompkins DM. Parasite spillback: a neglected concept in invasion ecology. Ecology. 2009;90: 2047–2056. doi: 10.1002/ecy.2446

18. Beckstead J, Meyer SE, Connolly BM, Huck MB, Street LE. Cheatgrass facilitates spillover of a seed bank pathogen onto native grass species. J Ecol. 2010;98: 168–177. doi: 10.1111/j.1365-2745.2009.01599.x

19. Dietrich R, Ploss K, Heil M. Growth responses and fitness costs after induction of pathogen resistance depend on environmental conditions. Plant, Cell Environ. 2005;28: 211–222. doi: 10.1111/j.1365-3040.2004.01265.x

20. Bradley DJ, Gilbert GS, Martiny JBH. Pathogens promote plant diversity through a compensatory response. Ecol Lett. 2008;11: 461–469. doi: 10.1111/j.1461-0248.2008.01162.x

21. Allan E, Van Ruijven J, Crawley MJ. Foliar fungal pathogens and grassland biodiversity. Ecology. 2010;91: 2572–2582. doi: 10.1890/09-0859.1

22. Stinson K, Kaufman S, Durbin L, Lowenstein F. Impacts of garlic mustard invasion on a forest understory community. Northeast Nat. 2007;14: 73–88. doi: 10.1656/1092-6194(2007)14[73:iogmio]2.0.co;2

23. Flory SL, Clay K. Invasive plant removal method determines native plant community responses. J Appl Ecol. 2009;46: 434–442. doi: 10.1111/j.1365-2664.2009.01610.x

24. Flory SL, Kleczewski N, Clay K. Ecological consequences of pathogen accumulation on an invasive grass. Ecosphere. 2011;2: art120. doi: 10.1890/ES11-00191.1

25. Gilbert GS, Webb CO. Phylogenetic signal in plant pathogen-host range. Proc Natl Acad Sci. 2007;104: 4979–4983. doi: 10.1073/pnas.0607968104

26. Malmstrom CM, Hughes CC, Newton LA, Stoner CJ. Virus infection in remnant native bunchgrasses from invaded California grasslands. New Phytol. 2005;168: 217–230. doi: 10.1111/j.1469-8137.2005.01479.x

27. Stricker KB, Harmon PF, Goss EM, Clay K, Luke Flory S. Emergence and accumulation of novel pathogens suppress an invasive species. Ecol Lett. 2016;19: 469–477. doi: 10.1111/ele.12583

28. Inouye B. Response surface experimental designs for investigating interspecific competition. Ecology. 2001;82: 2696–2706. doi: 10.1890/0012-9658(2001)082[2696:RSEDFI]2.0.CO;2

29. Hart SP, Freckleton RP, Levine JM. How to quantify competitive ability. J Ecol. 2018;106: 1902–1909. doi: 10.1111/1365-2745.12954

30. Fairbrothers DE, Gray JR. Microstegium vimineum (Trin.) A. Camus (Gramineae) in the United States. Bull Torrey Bot Club. 1972;99: 97. doi: 10.2307/2484205

31. Flory SL, Clay K. Non-native grass invasion suppresses forest succession. Oecologia. 2010;164: 1029–1038. doi: 10.1007/s00442-010-1697-y

32. Farr DF, Rossman AY. Fungal Databases. In: U.S. National Fungus Collections, ARS, USDA [Internet]. 2019 [cited 28 May 2020]. Available: https://nt.ars-grin.gov/fungaldatabases/

33. Drechsler C. Zonate eyespot of grasses caused by Helminthosporium giganteum. J Agric Res. 1928;37: 473–492. Available: https://naldc.nal.usda.gov/download/IND43967615/PDF

34. Lane B, Stricker KB, Adhikari A, Ascunce MS, Clay K, Flory SL, et al. Large-spored Drechslera gigantea is a Bipolaris species causing disease on the invasive grass Microstegium vimineum. Mycologia. 2020;00: 1–11. doi: 10.1080/00275514.2020.1781495

35. Kleczewski NM, Flory SL, Clay K. Variation in pathogenicity and host range of Bipolaris sp. causing leaf blight disease on the invasive grass Microstegium vimineum. Weed Sci. 2012;60: 486–493. doi: 10.1614/WS-D-l

36. Bürkner P-C. brms: An R package for Bayesian multilevel models using Stan. J Stat Softw. 2017;80: 1–28. doi: 10.18637/jss.v080.i01

37. Carpenter B, Gelman A, Hoffman MD, Lee D, Goodrich B, Betancourt M, et al. Stan: A probabilistic programming language. J Stat Softw. 2017;76. doi: 10.18637/jss.v076.i01

38. R Core Team. R: A language and environment for statistical computing. Vienna, Austria: R Foundation for Statistical Computing; 2018. Available: http://www.r-project.org/

39. Kay M. Tidy Data and Geoms for Bayesian Models. 2019. doi: 10.5281/zenodo.1308151

40. Wickham H, Averick M, Bryan J, Chang W, McGowan L, François R, et al. Welcome to the Tidyverse. J Open Source Softw. 2019;4: 1686. doi: 10.21105/joss.01686

41. Vloutoglou I, Kalogerakis SN. Effects of inoculum concentration, wetness duration and plant age on development of early blight (Alternaria solani) and on shedding of leaves in tomato plants. Plant Pathol. 2000;49: 339–345. doi: 10.1046/j.1365-3059.2000.00462.x

42. Hetherington SD, Smith HE, Scanes MG, Auld BA. Effects of some environmental conditions on the effectiveness of Drechslera avenacea (Curtis ex Cooke) Shoem.: A potential bioherbicidal organism for Avena fatua L. Biol Control. 2002;24: 103–109. doi: 10.1016/S1049-9644(02)00020-8

43. Chen L, Zhou J, Zeng T, Miao Y, Mei L, Yao G, et al. Quantifying the sharing of foliar fungal pathogens by the invasive plant Ageratina adenophora and its neighbours. New Phytol. 2020;3. doi: 10.1111/nph.16624

44. Johnson NC, Graham JH, Smith FA. Functioning of mycorrhizal associations along the mutualism-parasitism continuum. New Phytol. 1997;135: 575–585. doi: 10.1046/j.1469-8137.1997.00729.x

45. Klironomos JN. Feedback with soil biota contributes to plants rarity and invasiveness in communities. Nature. 2002;417: 67–69. doi: 10.1038/417067a

46. Mangan SA, Schnitzer SA, Herre EA, MacK KML, Valencia MC, Sanchez EI, et al. Negative plant-soil feedback predicts tree-species relative abundance in a tropical forest. Nature. 2010;466: 752–755. doi: 10.1038/nature09273

47. Dostál P, Müllerová J, Pyšek P, Pergl J, Klinerová T. The impact of an invasive plant changes over time. Ecol Lett. 2013;16: 1277–1284. doi: 10.1111/ele.12166

48. Levine JM, Adler PB, Yelenik SG. A meta-analysis of biotic resistance to exotic plant invasions. Ecol Lett. 2004;7: 975–989. doi: 10.1111/j.1461-0248.2004.00657.x

49. Orrock JL, Dutra HP, Marquis RJ, Barber N. Apparent competition and native consumers exacerbate the strong competitive effect of an exotic plant species. Ecology. 2015;96: 1052–1061. doi: 10.1890/14-0732.1

